# Fast Loading and Transformation of Large-Volume Bioimaging Data Consistently Approaching the I/O Peak Speed

**DOI:** 10.64898/2025.12.10.693603

**Authors:** Lin Cai, Xuzhong Qu, Hang Zhou, Ning Li, Xianyu Gou, Shiwei Li, Jian Huang, Shasha Guo, Xiuli Liu, Xiaohua Lv, Tingwei Quan, Shaoqun Zeng

## Abstract

High-resolution large-volume biological imaging techniques are now widely used in biological research, but inefficiencies in data processing and visualization persist due to bottlenecks in loading/saving pipelines and limitations of conventional pyramid formats. To address these issues, we have decoupled data loading process into reading from disk and in-memory decompression, while data saving has been separated into in-memory compression and writing to disk. This approach allows us to dedicate more CPU cores to decompression and compression tasks, optimizing resource allocation and maximizing disk I/O throughput. As a result, both loading and saving processes operate near the disk’s peak data transfer capacity. Further, we integrated a video stream-inspired format into the pyramid structure, allowing direct access to regions of interest (ROIs) without loading extraneous data into RAM, reducing memory overhead. Compared to the latest methods, our approach improves pyramid data transformation efficiency by at least 6× and dramatically accelerates downstream tasks like visualization and deep learning inference.

**Teaser:** Novel bioimaging data processing pipeline achieves near-peak I/O throughput via decoupled architecture and video-formatting.

## Introduction

Recent advancements in high-throughput optical imaging techniques, such as light-sheet imaging and micro-optical sectioning tomography, have made it possible to observe large-scale tissues with sub-cellular resolution(*1-5*). These developments greatly enhance our understanding of tissue structure and function. Simultaneously, the datasets generated from a single sample acquisition can easily reach terabyte (TB) in size, leading to an explosive growth in imaging data. Managing, visualizing, and analyzing these TB-sized images is crucial for mining valuable information and constructing biophysical models at various scales(*6-8*). These data processing tasks require loading data from external storage to random access memory (Data loading), followed by saving analytical results back to external storage (Data saving). Therefore, the efficiency of data loading and saving has a direct impact on the speed at which large-volume images are processed.

To accelerate massive data processing, numerous studies have focused on pyramid data transformation(*9-17*), which converts an image sequence into a pyramid structure. This structure stores smaller volume of data at multiple resolutions and enables efficient loading of specific data blocks within the large-volume image based on user requirements. The Hierarchical Data Format version 5 (HDF5) serves as a flexible and widely supported format for this purpose(*12*). Based on HDF5, the BigDataviewer optimizes random access to large datasets and allows for rapid browsing of regions of interest(*13*). Terafly(*14*) and Tdat(*11*) have optimized the entire workflow for pyramid data transformation, enabling them to handle petabyte-sized image volumes. N5 addresses the inefficiencies associated with HDF5 in loading and saving large-volume data by organizing a large dataset into multiple files to facilitate the use of parallel techniques(*15*). Zarr(*16, 17*) offer an effective solution for high-frequency data access by large numbers of users, facilitating powerful tools for bio-image visualization and analysis(*18-21*). Additionally, several approaches have been developed to enhance data loading and saving processes. OME-Tiff(*22*) combines the benefits of OME-XML and TIFF formats, allowing each TIFF file to include rich metadata defined by OME-XML, which broadens its applicability. Tifffile(*23*) takes advantage of the memory mapping technique to process large images. Cpp-Tiff(*24*) leverages the OpenMP(*25*) framework to parallelize LibTiff’s code, improving both loading and saving speeds.

Despite significant progress in data loading/saving and pyramid data transformation, two issues remain unresolved. In the current data loading mode, data reading and decompression are handled together, leading to the allocation of the same Central Processing Unit (CPU) resources (equal threads) for both tasks. This is an inefficient allocation, as data decompression is far more time-consuming than data reading. A similar issue occurs in data saving, where data compression and writing are combined. These inefficiencies significantly reduce overall data processing performance. In addition, the chunk format used in pyramids is closely tied to Tiff, HDF5 or Zarr which requires loading large amounts of redundant data surrounding the region of interest (ROI) into Random Access Memory (RAM) for visualization and analysis, which increases the burden on data loading and processing operations.

In this work, we redesigned the data loading and saving processes in LibTiff. For data loading, we separated reading and decompression to allocate more CPU threads to decompression. Similarly, during data saving, compression and writing were decoupled to enhance efficiency. Additionally, we introduce a new pyramid data structure, the Biomedical Video Stream (Bio-VS) format, which is based on the video format used in FFmpeg rather than Tiff, HDF5 or Zarr. This new format supports direct access to any specific frame data within chunks and achieves a higher compression ratio. These two design improvements significantly enhance the efficiency of data transformation, visualization, and deep learning inference. To facilitate usage, we developed Decoupled-Tiff (Dec-Tiff), a Python library for rapid loading and saving of data in both Tiff format and Bio-VS formats. We also created BioimageVision, a software tool that enables the direct pyramid data transformation and visualization of bio-images.

## Results

### Data loading and saving

Data loading and saving are crucial steps in the process of visualizing and analyzing large-volume images (**Figure 1a**). Initially, images stored on disk must be loaded into RAM (Data loading), where they are reorganized into a pyramid structure. This pyramid data comprises multi-resolution chunks that require compression and subsequent storage back to disk (Data saving). Users can efficiently access ROI from the pyramid data for further analysis. LibTiff serves as a fundamental library tool for data loading and saving, with variants such as OME-Tiff, Tifffile, and Cpp-Tiff significantly boosting efficiency in these tasks. In these library tools, during the data loading process, data reading and decompression are combined and treated as a single operation (**Middle panel in Figure 1a**). In contrast, our library tool, Dec-Tiff, redesigns certain components of LibTiff. In Dec-Tiff, data reading and decompression are decoupled during data loading (**Light green panel in Figure 1a**). By using this decoupling strategy, we can assign a single CPU thread for data reading and multiple CPU threads for data compression. Essentially, this design allows us to approximate the time cost associated with data loading as almost equivalent to the duration spent on reading the data alone, rather than summing both reading and decompression times cumulatively (**Figure 1b**). This design is based on the fact that the time required for data reading is significantly less than that needed for decompression (**Fig. S1**). The same operations are also applied to data saving in Dec-Tiff.

**Figure 1.**
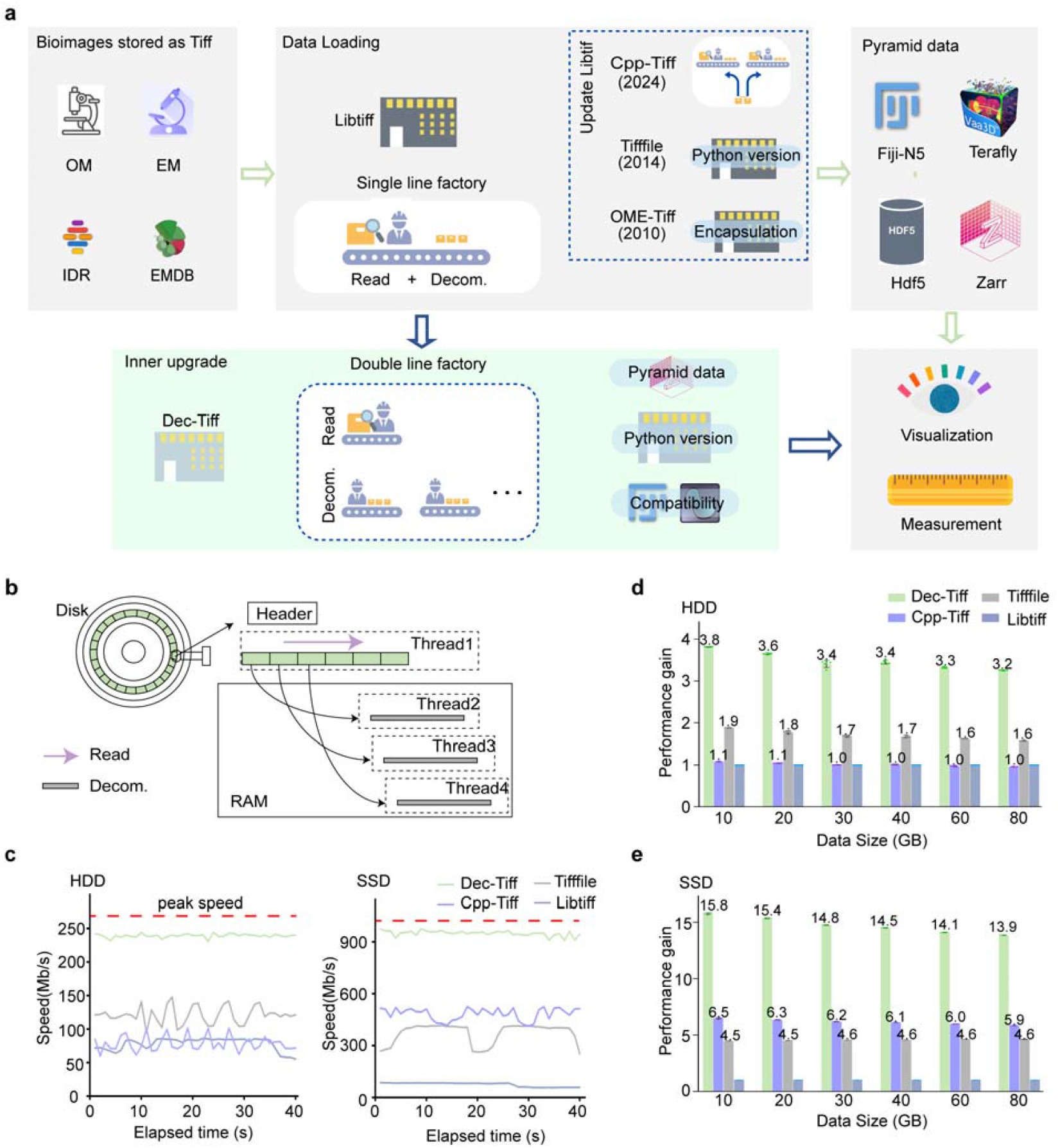
Performance comparison of image loading modes with and without decoupling strategy. **(a)** The overall workflow for handling large-volume imaging data. (i) Utilizing an image loading-saving library for loading a TIFF image sequence. (ii) Transforming an image sequence into pyramid data. (iii) Using the pyramid data format for fast localization of regions of interest and conversion across different resolutions, visualization and quantitative analysis of large-volume data are achieved. During the data loading process, data reading and decompression have been decoupled through a redesign of relevant code sections in LibTiff. This redesign has led to the creation of a new data loading library named Dec-Tiff, represented by the light green region. **(b)** The process of loading images with Dec-Tiff involves several steps: first, extracting the header file information; second, identifying and locating the necessary objects for reading; third, performing serial data reading; and fourth, executing parallel data decompression. **(c)** Comparison of image loading speeds on hard disk drive (HDD) and solid-state drive (SSD) among the decoupled mode (Dec-Tiff), Cpp-Tiff, Tifffile, and LibTiff. **(d)** Performance improvements of Dec-Tiff over Cpp-Tiff and Tifffile in terms of loading 3D image stacks of uint16 type, with dimensions 650×500×256 (x,y,z), on an HDD. Each test was conducted independently five times on an i9-13900KF CPU with 8 cores enabled, and the data are presented as mean ± standard deviation (std). **(e)** Similarly, performance improvements of Dec-Tiff over Cpp-Tiff and Tifffile for loading the same 3D image stacks on an SSD. Each test was also independently run five times on the same CPU configuration, and the results are reported as mean ± std.

We quantify the efficiency of data loading to demonstrate the advantages of this decoupling strategy. The test data is a group of volume images, each of which has the size of 650×500×256 and is LZW compressed. They are stored on a hard disk drive (HDD) and a solid state drive (SSD), respectively. In data loading or saving, the computer automatically presents their speeds which are used to measure the I/O resources utilization performance. Dec-Tiff achieves an average speed of 247 MB (Megabyte) per second in data loading on HHD (**Left panel in Figure 1c**), almost consistently close to the peak I/O speed of the HDD (270 MB/s). On SSD, the loading speed is also consistently close to its peak I/O speed (**Right panel in Figure 1c**). This loading efficiency is far superior to that of Cpp-Tiff, Tifffile and LibTiff.

We compared the performance of Dec-Tiff with several other library tools in data loading and saving. The test data was identical to that in **Figure 1c** and its size was adjusted by modifying the number of image blocks. For HDD, Dec-Tiff demonstrated a speed that was 3 times faster than LibTiff and 2 times faster than Tifffile (**Figure 1d**). On SSD, Dec-Tiff outperformed LibTiff by 14 times and was 2 times faster than Cpp-Tiff, which is specifically optimized for SSD data loading (**Figure 1e**). Furthermore, we compare the performance of these tools on different number of CPU cores and data block with different number of frame (**Fig. S2**). During the data loading process, SSD leverage electrical addressing to swiftly locate required data, fulfilling the requirements of multiple CPU cores and rendering the parallel mode effective. However, due to the slower addressing speed of HDD, this parallel mode does not enhance performance (**Figure 1c**), limiting its applicability. Dec-Tiff successfully addresses this unresolved issue and significantly enhances data loading efficiency on HDD. For data saving, Dec-Tiff was also 2-3 times faster than other tools on both HDD and SSD (**Fig. S3**), performing equally well as it did during data loading.

### Data transformation

We apply our data loading and saving method for pyramid data transformation. In the process of generating pyramid format data, the common steps are briefly described as follows: read in the two-dimensional image and decompress it into a two-dimensional array; perform multi-scale sampling on these two-dimensional arrays; stack them in order to form data blocks in a pyramid structure; compress the data blocks and write them to the disk. The main difference between our workflow and others is that we decouple the processes of reading and decompression, as well as compression and writing (**Figure 2a**). This strategy decomposes the entire workflow into more steps, and allows for more flexible allocation of CPU threads. The CPU threads are automatically assigned into each stage of our data transformation to prevent data flux blocking (**Materials and Methods**). In our data transformation, a number of local images in an image sequence can be localized by extracting the header file information, and then converted into a portion of pyramid data. The chunks in the pyramid data are the video stream format (Bio-VS) and the data in z-direction is not partitioned (**Figure 2a**). Bio-VS is more concise and effective compared to the current pyramidal data structure (**Fig. S4**). The video stream format supports direct access to arbitrary frame data. Due to this characteristic, when manipulating the specified images across multiple chunks, these images can be directly accessed without loading the associated chunks into RAM (**Fig. S4**).

**Figure 2.**
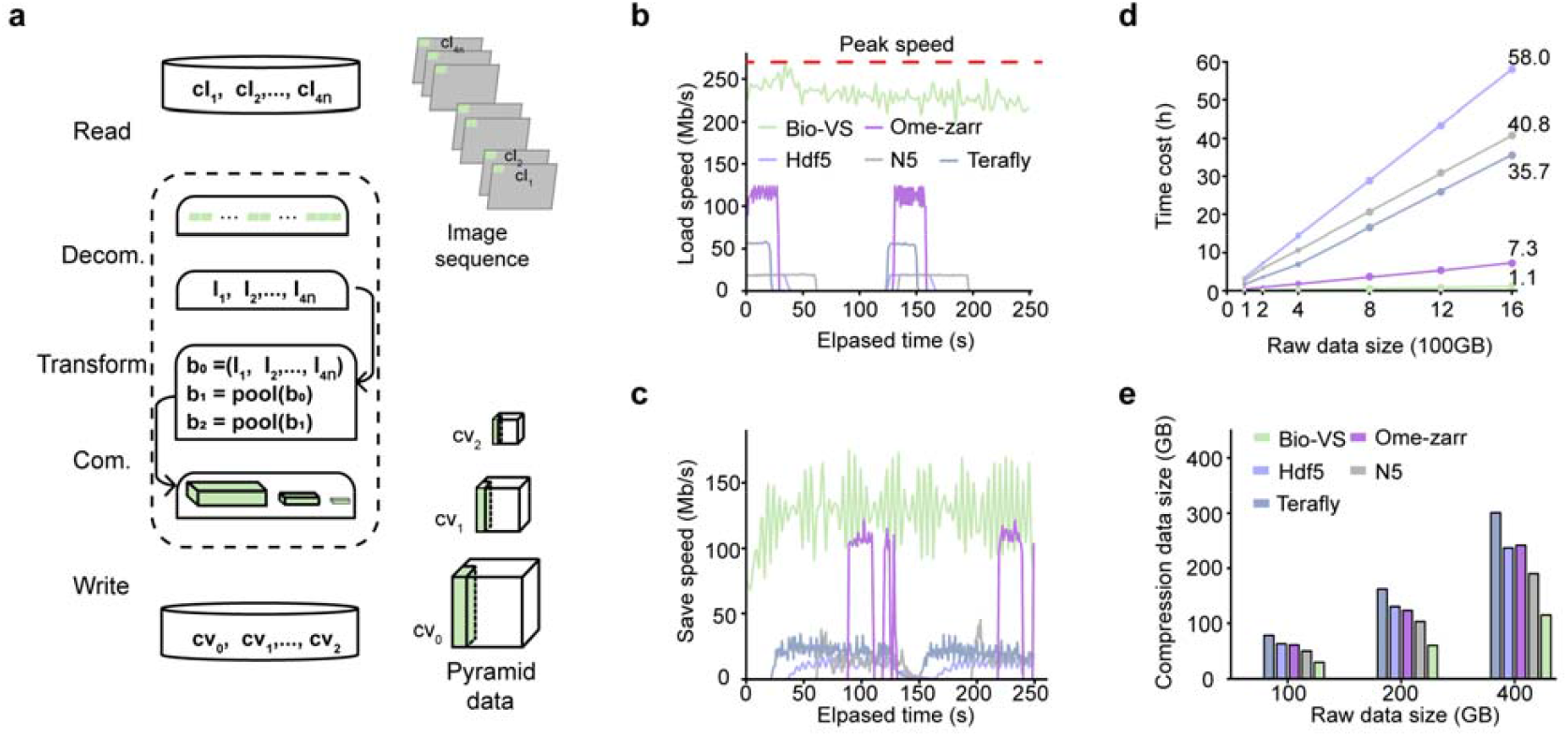
Performance in creating pyramid image structures. (**a**) The schematic overview of the proposed data transformation workflow converting an image sequence into the Bio-VS format. The workflow includes raw data reading, lossless decompression, data transformation, compression and structured output writing. (**b**) Comparisons of data loading speeds over time between the Bio-VS framework and conventional formats (Ome-zarr, Hdf5, N5, Terafly). (**c**) Corresponding evaluation of data saving speed measured during the same experimental interval as (**b**). (**d**) Quantitative comparison of computation time required for pyramid data structure generation in Bio-VS versus competing formats. (**e**) The compressed data sizes of different pyramid data formats originating from uncompressed baseline data, collected from the fMOST system, with the sizes of 8192×8192×1600, 8192×8192×3200, 16384×16384×1600 respectively.

Our data transformation consistently approaches the I/O peak speed of an HDD during data loading processes (**Figure 2b**), with all CPU cores being fully utilized throughout the entire workflow. The test data are a set of volumetric images collected from collected from Fluorescence micro-optical sectioning tomography system (fMOST)(*26*). Each volumetric image consists of 8-bit two-dimensional Tiff image sequences, with each image measuring 16384×16384 in size. It is worth noting that the data saving speed is approximately 130 MB/s lower than the peak I/O speed. This is reasonable since the transformed data is saved in a video stream format that inherently has a higher decompression ratio (**Figure 2c**). These results indicate that there is no blockage in the data flow throughout the entire data transformation process. We evaluated the performance of several freely available tools for data transformation. In a single loop, both the data saving and loading speeds were below 40 MB/s. Furthermore, when transforming a set of volumetric images ranging in size from 100 GB to 1600 GB stored on an HDD, our method achieved at least a 6× reduction in time compared to OME-Zarr and approximately a 30× reduction compared to N5, HDF5, and Terafly (**Figure 2d**). For these tools, our test experiments utilized their default modes, which fully consider parallel processing. Additionally, our transformed volumetric images are lossless compression (**Fig. S5**) and exhibit the highest compression ratio (**Figure 2e**), making them advantageous for data storage and transmission.

We further analyzed the performance characteristics of Bio-VS and OME-Zarr on fMOST datasets with varying properties, comparing both pyramid data generation time and compression efficiency. Experimental results demonstrate that, compared to the OME-Zarr, the Bio-VS achieves a 5-9 times improvement in pyramid construction speed, while the size of compressed data can be reduced by up to 50% (**Table. S1**).

### Visualization of volume datasets

We have also developed a software called BioimageVision for data visualization and transformation. Our data visualization process employs a loading strategy similar to that used for TIFF images, which separates data reading and decompression. This approach leverages multiple CPU threads to decompress the data in parallel, significantly speeding up the process. Additionally, the Bio-VS format enables direct access to specified data within the pyramidal structure, minimizing the loading of redundant data as much as possible. These two optimizations collectively significantly reduce response times during visualization tasks.

To demonstrate the advantages of Bio-VS, we evaluated 16.9 TB whole-mouse-brain image datasets acquired using fMOST(*26*) technology. These datasets were converted into both Bio-VS format and the conventional OME-Zarr pyramid structure for comparative analysis. We compared the visualization efficiency of these two formats for both local volumes of interest and 2D data slices, using response time, defined as the total duration required for data readout and decompression, as the metric to measure visualization efficiency. In data block visualization (**Figures 3a-c**), Bio-VS demonstrated a twofold improvement in response time compared to OME-Zarr (**Figure 3d**). This enhancement is primarily attributed to Bio-VS’s optimized data loading mechanism, which decouples data reading and decompression processes. When visualizing 2D slices (**Figures 3e-g**), Bio-VS enables direct and precise targeting of individual slices, whereas OME-Zarr’s structure requires loading of the related data chunks, including redundant adjacent slices. As a result, Bio-VS exhibits a linear scaling of response time with increasing slice size, while OME-Zarr shows a nonlinear due to the expanded loading of redundant data (**Figure 3h**). For large-scale slice visualization, Bio-VS provides at least a 12-fold improvement in response time (**Figure 3h**), enabling real-time visualization of entire mouse brain slices. These results demonstrated that Bio-VS outperforms OME-Zarr in both block and slice-based visualization tasks, offering significant improvements in speed and scalability.

**Figure 3.**
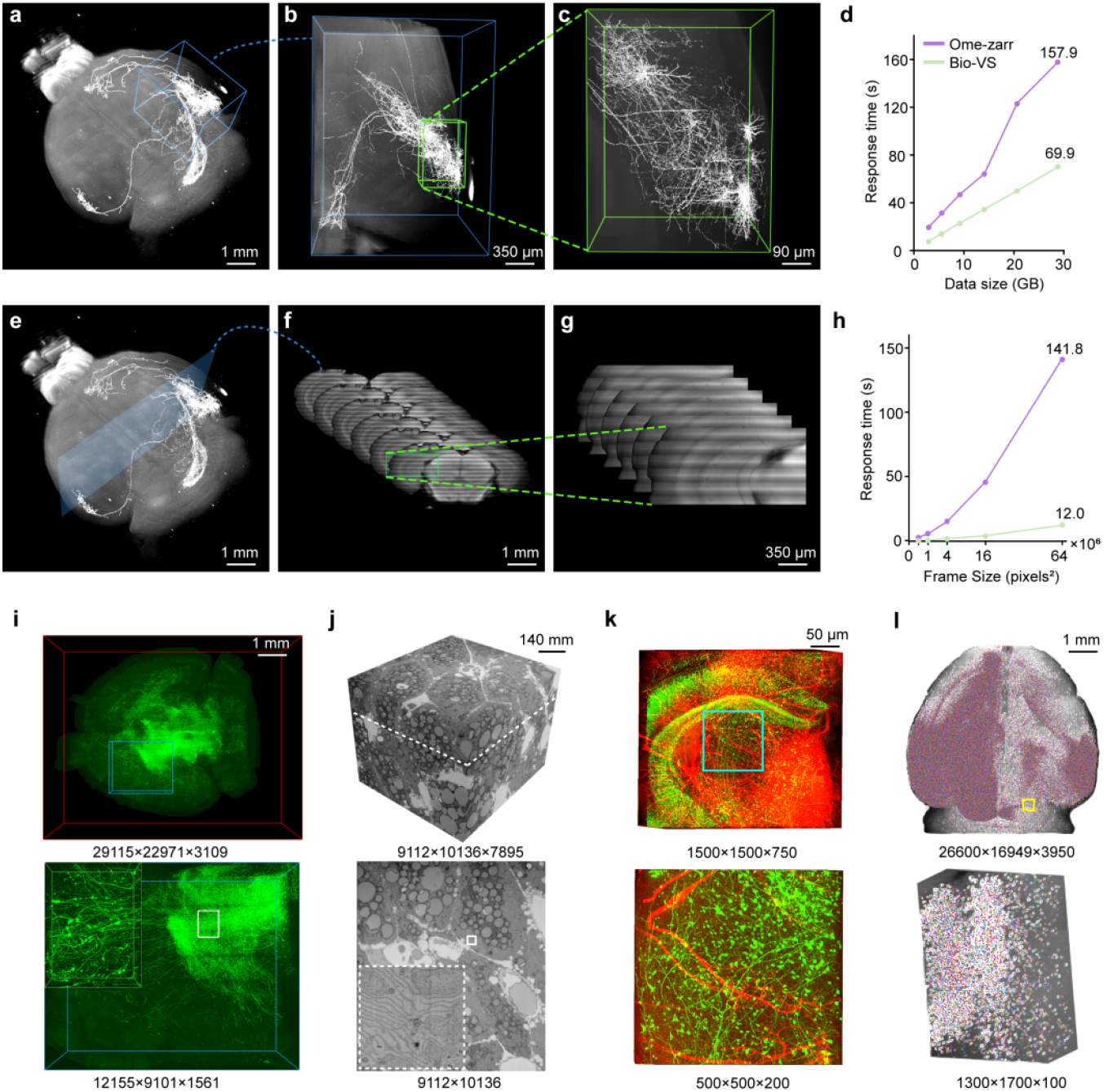
The performance of Bio-VS data visualization and its response speed. (**a-c**) Visualization of different multi-resolution data blocks in Bio-VS data. This Bio-VS data is originated the whole mouse brain fMOST imaging data. (**d**) The comparison of visualization response speeds between the Bio-VS and Ome-zarr formats across data blocks of varying sizes. The raw imaging data used in (**a**) is also transformed into Ome-zarr data. The same data blocks in these two pyramid datasets are visualized and the corresponding visualization response times are calculated. (**e-g**) Visualization of multi-resolution data slices in Bio-VS data used in (**a**). (**h**) Visualization response times concerning 2D slices in the formats of Bio-VS (green) and Ome-zarr (purple). The testing datasets comprise 2D slices varying from 500×500 to 8000×8000 pixels. (**i-l**) Visualization of 3D images collected with light-sheet microscopy (LSM), volumetric images collected with scanning electron microscopy (SEM), two-channel 3D images collected with fMOST and the cell positions mapped within the volume data.

BioimageVision enables efficient processing and visualization of large-scale bioimaging datasets. Here, we demonstrate its applicability to terabyte-scale datasets acquired via light-sheet microscopy and scanning electron microscopy (**Figures 3i&j**). Our data transformation pipeline achieves a throughput of ∼1.3 terabytes per hour, underscoring optimized utilization of HDD I/O resources across diverse imaging modalities and data sizes. Post-transformation, volumetric datasets are rendered interactively at a sustained rate of 512×512×512 voxels/second, facilitating seamless navigation across ROIs. BioimageVision offers a diverse range of visualization capabilities, including the visualization of multi-channel volumetric images (**Figure 3k**), as well as cell positions (**Figure 3l**) and reconstructed neurons (**Fig. S6**), which fulfills the fundamental needs for imaging data visualization.

### Bio-VS data compatibility with open-source software

The Bio-VS data can be exported to popular open-source software platforms such as Napari(*27*) and Fiji(*28*) to support quantitative analysis workflows (**Figure 4a**). By installing the Bio-VS plugin in Napari, users can incorporate Bio-VS data into their analytical workflows (**Figure 4b**). This integration allows for efficient loading and visualization of large-scale image datasets, and the flexibility to choose ROI for targeted quantitative measurements.

**Figure 4.**
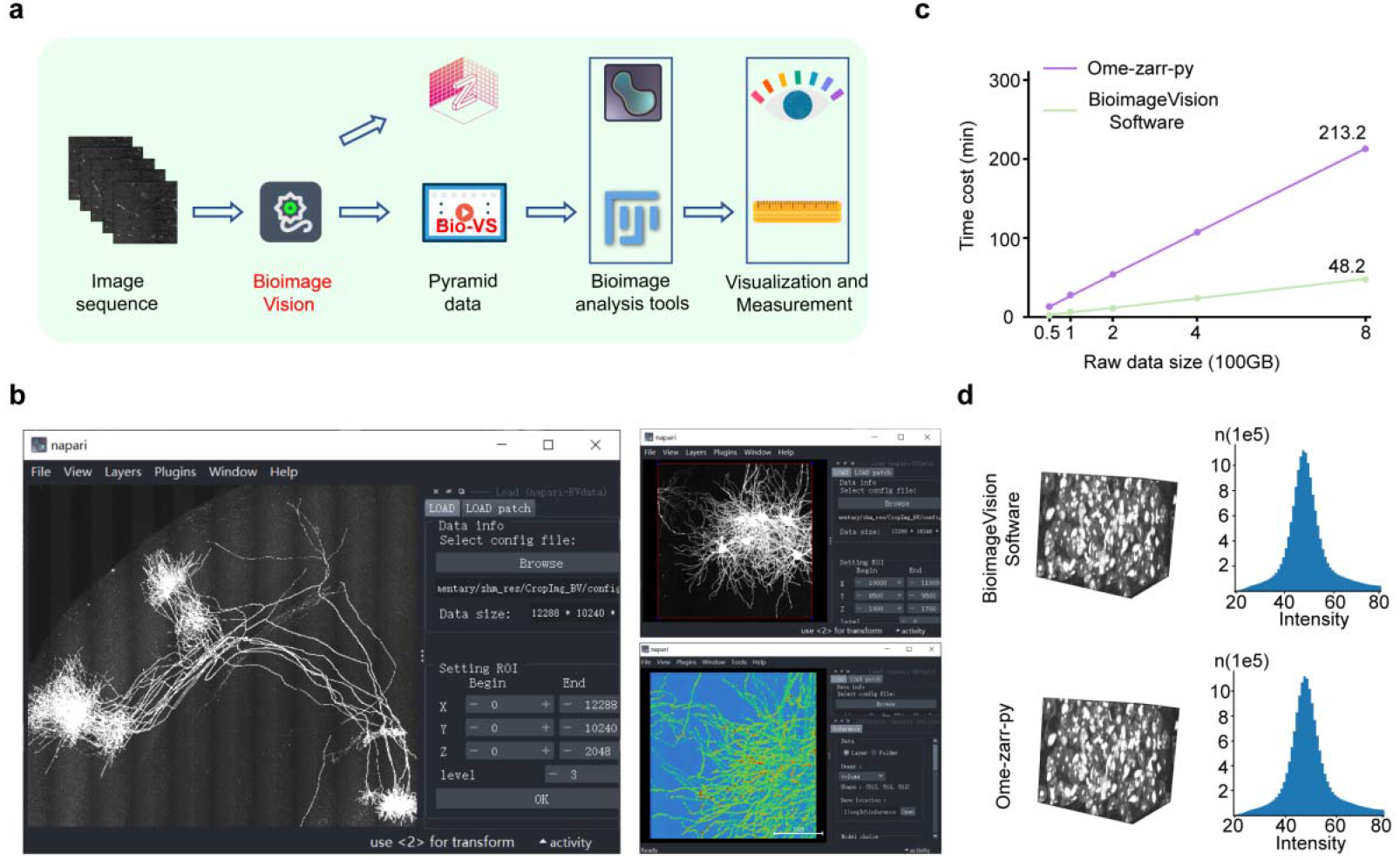
Compatibility of Bio-VS format. (**a**) The overall workflow for ensuring the compatibility of the Bio-VS format with existing mainstream software. Image sequences can be converted into Bio-VS format using BioimageVision software, which also offers rapid conversion to the Ome-zarr format. With our plugins or scripts, the Bio-VS format can be seamlessly loaded into tools like Napari, Fiji and others for further visualization and measurement. (**b**) Bio-VS data can be loaded into Napari, enabling interactive visualization, manipulation, and processing of the data within a user-friendly interface. (**c**) BioimageVision software significantly accelerates Ome-zarr format generation from image sequences compared to the official Ome-zarr Python interface. (**d**) Statistical analysis confirms the precision of Ome-zarr data generated by BioimageVision. Histograms of data blocks produced by both BioimageVision and the official Ome-zarr Python interface were identical.

In addition to the integration with Napari, we developed a Python script that automates Fiji execution via PyImageJ, creating a streamlined analysis pipeline. This script enables direct import of Bio-VS datasets into Fiji, where users can select and analyze specific regions of interest (ROIs) with minimal manual intervention (**Fig. S7**). By leveraging Fiji’s built-in image processing tool, including segmentation, colocalization, and intensity measurement functions, users can perform comprehensive quantitative analyses on Bio-VS data. The workflow ensures end-to-end automation, from data transfer to ROI-focused computation, while maintaining access to Fiji’s established analytical plugins.

OME-Zarr is a widely adopted and supported format for managing large-scale imaging datasets due to its efficient compression and metadata handling. BioimageVision also achieves the conversion of image sequences into OME-Zarr format using optimized data loading and saving method (**Figure 4a**). However, inherent limitations of the OME-Zarr specification, which prioritize metadata consistency over raw I/O speed, mean that hard drive input/output (HDD I/O) performance is not fully maximized during this process. Despite this constraint, BioimageVision achieves a conversion rate of 1 TB per hour, marking a 4× speedup compared to the Ome-Zarr-py library when tested on fMOST and light-sheet datasets (**Figure 4c**). In addition, the OME-Zarr outputs generated by BioimageVision are validated to be identical to those produced by Ome-Zarr-py **(Figure 4d**), ensuring compatibility with existing analysis pipelines while enhancing throughput.

### The enhancement in the inference speed of deep networks

As the computational power of GPUs continues to expand, deep learning models are increasingly tasked with analyzing vast biomedical images that often reach sizes of hundreds of gigabytes or even terabytes. However, data loading has emerged as a critical bottleneck in this process. Traditional methods struggle to transfer such large volumes of data swiftly from storage drives (HDD/SSD) into GPU memory, resulting in under-utilization of powerful multi-GPU systems.

To address this challenge, we integrated our data loading into the network inference pipeline. The pipeline facilitates rapid in-memory loading of Bio-VS formatted data, efficient distribution across multiple GPUs, and seamless integration into the neural network (**Figure 5a**). We demonstrate this methodology using a whole mouse brain image comprising dimensions of 26,600 × 15,888 × 8,597 voxels. The Bio-VS data structure organizes the image into a pyramid format; the highest-resolution layer (base blocks) contains chunks, each with 512×512×512. These chunks are processed by the 3D U-Net(*29*) for the segmentation of whole brain images.

**Figure 5.**
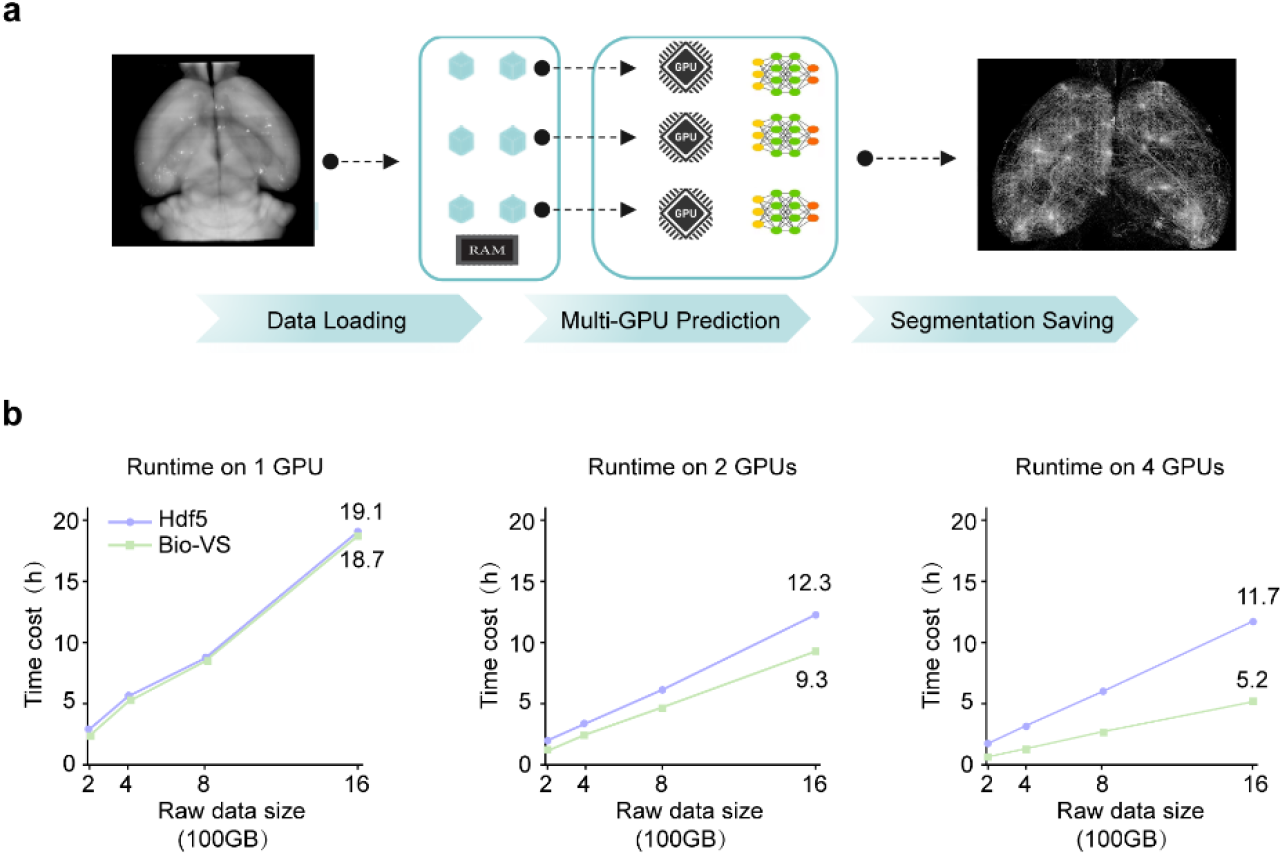
The application of Bio-VS data format in deep learning networks for whole-brain scale neural image segmentation. (**a**) The workflow of Bio-VS applied in the segmentation task. In this workflow, Bio-VS data is generated from the whole mouse brain imaging data, and the image blocks in Bio-VS structure are loaded into RAM and then inputted multi GPUs for segmentation network inference. (**b**) Comparisons of the time cost associated with the deep learning network inference with Bio-VS and Hdf5 data inputs. For Bio-VS, the time required for network inference decreases proportionally with the number of GPUs (RTX 3090) utilized. In contrast, for Hdf5, when the number of GPUs increases from 2 to 4, there is almost no change in the network inference time.

We quantitatively assessed how the efficiency of the network inference pipeline scales with the number of GPUs (RTX 3090). When utilizing two GPUs versus four GPUs in the pipeline, the total inference time decreased proportionally: halving with two GPUs and quartering with four GPUs, relative to single-GPU performance (**Light green curves in Figure 5b**). This linear reduction indicates that the Bio-VS data structure’s loading efficiency effectively matches with increasing GPU resources. In contrast, pipelines using conventional HDF5 data loading exhibited minimal improvement in inference time when scaling from two to four GPUs (**Light blue curves in Figure 5b**), which suggests that HDF5’s data loading speed becomes a bottleneck, limiting the pipeline’s ability to leverage additional GPUs. Collectively, these results demonstrate that integrating our data loading strategy with the Bio-VS structure significantly enhances inference efficiency for large-scale image analysis, enabling scalability with GPU resources.

### The use of the developed tools

The BioimageVision software is focused on data transformation and visualization capabilities. In terms of data transformation, it facilitates the conversion of LZW compressed TIFF images into the Bio-VS format. To optimize performance, we recommend distributing Bio-VS data and source imagery across two separate hard disk drives (HDDs). One is dedicated to data loading and the other for saving to mitigate I/O bottlenecks and ensure uninterrupted data throughput. Furthermore, the software also converts LZW compressed TIFF files to the Zarr format, utilizing our efficient data loading and saving method that significantly enhance processing speeds.

The BioimageVision software enables users to export ROIs as TIFF images for downstream quantitative analysis tasks, including object segmentation, localization, and skeletonization. Results from these analyses which are stored in either SWC format(*30*) (for neuron morphology measurement) or TIFF images, can be visualized alongside their corresponding ROIs within the Bio-VS datasets. This workflow addresses BioimageVision’s limitation in directly annotating and analyzing 3D volumetric data. Additionally, Bio-VS data compatibility with tools like Fiji and Napari expands access to advanced image analysis algorithms.

We present a Python library function called Dec-Tiff, which is built upon our data loading and saving methods, and operates independently of the BioimageVision software. Dec-Tiff allows users to efficiently load Bio-VS format data into memory and convert it into arrays for subsequent analysis. Furthermore, Dec-Tiff utilizes our optimized data loading mode to enable rapid reading and writing of 2D 8/16-bit TIFF images. We demonstrate that Dec-Tiff effectively loads Bio-VS data for deep network inference. Overall, Dec-Tiff offers a relatively streamlined approach to enhance the applicability of our data loading and saving methods for large-scale image visualization and analysis.

## Discussion

We have developed image loading and saving methods that fully leverage the I/O resources of both HDD and SSD. In conjunction with the video stream format, these methods efficiently transform image sequences into pyramid-shaped data structures. Building on these two innovations, we achieve rapid visualization of large-scale 3D images and the enhancement in the inference speed of deep networks.

In our data transformation process, the throughput of the data flow consistently approaches the maximum value permissible by the computer. The crucial aspect is that data decompression and compression are decoupled from data reading and writing. Without this decoupling strategy, direct parallel data loading or saving fails to achieve this high throughput, as illustrated in Figure 2b and 2c. The underlying reason is that parallel threads access the disk simultaneously, leading to severe I/O contention. In addition to this decoupling strategy, we also fully utilize CPU cores and automatically allocate them at each stage of the data transformation process to ensure minimal data flow bottlenecks. Therefore, our data transformation demands relatively high standards for computer CPU performance; however, the requirement for random access memory (RAM) capacity remains comparatively low. These needs contrast with tools like OME-Zarr-py and Terafly, which both necessitate substantial RAM capacity for loading data blocks.

We have decoupled the processes of reading and decompressing Bio-VS data. This decoupling enables us to allocate more CPU cores specifically to the task of Bio-VS data decompression, resulting in rapid loading of the data into RAM. This, in turn, significantly enhances the speed of deep network inference and data visualization. Furthermore, our decoupling approach also boosts the generation speed of OME-Zarr files. The results demonstrate that our decoupling strategy can be effectively utilized in the analysis of large-scale volume datasets, irrespective of their formats. In terms of data visualization and deep network inference, our current data loading processes do not achieve peak I/O speeds. This limitation arises because optimization efforts have focused solely on data loading rather than addressing the entire processing pipeline. Ideally, both the speed for data visualization and the inference speed for deep networks should approach I/O peak speeds of HHD. Achieving this objective necessitates an appropriate allocation of computational resources across all processes within the analysis pipeline, in conjunction with our strategy for efficient data loading and saving. These operations are similar to those employed in our data transformation whose speed is close to I/O peak speeds of HHD.

A diverse range of image formats and compression algorithms exists, leading to numerous compressed image formats. Our data loading and saving methods are built upon the LibTiff library framework. Currently, these methods are specifically optimized for 8/16-bit TIFF 2D images using LZW compression. However, substantial future work is required to expand these capabilities and support broader application scenarios.

## Materials and Methods

### The data loading and saving

Given that image decompression incurs a greater time cost than image reading, and recognizing that disk-based image reading is inherently sequential, we adopt a parallelized resource allocation strategy. Specifically, we designate a single CPU thread for image reading tasks, while reserving multiple threads for decompression operations. During image readout, the system first parses the header metadata to determine the image’s disk location, followed by loading the image into RAM. Each image comprises numerous independently decompressible units, which, once resident in memory, activate the decompression cores. The resulting decompressed data is systematically stored in a preallocated RAM buffer.

For image saving operations, a preprocessing step establishes compression threads. These threads competitively utilize available CPU cores to compress image units, which are then appended to an ordered output queue. Concurrently, a single CPU thread manages the sequential writing of compressed units to persistent storage media (HDD/SSD), ensuring temporal coherence between processing and I/O stages. This architecture optimizes throughput by decoupling resource-intensive decompression/compression tasks from the inherently serial disk I/O operations.

Our decoupling strategy is compatible with all data formats supported by Libtiff. However, it is important to note that when decoupling a single image within a batch of images, whether during loading or saving, our decoupling strategy may result in speeds that approach the I/O limits of the disks. In addition, regarding the decoupling operation for image subregions, at present, it is capable of supporting the TIFF data format.

### The workflow of data transformation

The BioimageVision data transformation pipeline is structured as a five-phase workflow: data reading, decompression, transformation, compression, and saving. Within this framework, reading and saving stages operate sequentially, while the intermediate three phases execute in parallel through multi-core CPU utilization. To ensure optimal throughput, the system balances processed data volumes across all stages, with this equilibrium directly governing CPU core allocation per phase.

BioimageVision employs automated detection of total available CPU cores. To mitigate I/O contention, both reading and output operations are assigned a single dedicated CPU thread. The remaining threads are dynamically partitioned among decompression, transformation, and compression stages, operating in a competitive scheduling model to maximize resource utilization. The allocation algorithm prioritizes resource-intensive phases, allocating proportional computational power based on workload demands. Empirical testing demonstrates that moderately provisioned systems achieve reading rates exceeding 220MB/s, validating the architecture’s efficiency in balancing I/O and computational loads.

### The brief description of BioimageVision

This software is engineered using a hybrid architecture that integrates Python and C++. Python serves as the primary interface layer, ensuring user-friendly interaction and workflow orchestration, while C++ handles performance-critical computations, memory management, and computational resource allocation. The platform’s core capabilities encompass: 1) data transformation pipelines for pyramid data format conversion; 2) multi-resolution visualization of large datasets, supporting interactive exploration across scales; 3) advanced volumetric rendering, including maximum intensity projection (MIP) generation. Additionally, a Python API is provided to facilitate the conversion of BV formatted data into TIFF stacks, enabling seamless integration with downstream applications such as deep learning model training. The software is compatible with Windows 10 (64-bit) environments. For detailed system requirements, installation instructions, and usage guidelines, please refer to the accompanying User Guide.

### Test images description

The test data primarily comprises three types of datasets: light-sheet microscopy (LSM) data, electron microscopy (EM) data, and Fluorescence micro-optical sectioning tomography (fMOST) data. The LSM data, which has a total size of 3.78 TB, is described in detail in reference(*21*). The EM data, which has the size of 2.1 TB, is freely accessible, with a comprehensive description provided in reference(*31*). The fMOST data has a size of 16.07 TB, and its details can be found in reference(*26*). Both the fMOST and LSM datasets are predominantly utilized to evaluate the performance of data transformation and visualization techniques. The electron microscopy data serves as an illustrative example demonstrating that our method can be applied to any volumetric data formatted in Tiff.

### Performance evaluation

The compression ratio is defined as the quotient of the size of uncompressed data divided by the size of compressed data. Visualization response time refers to the total duration required for both data reading and decompression. Data transformation processes were carried out through software-based operations. To evaluate the efficiency of data transformation, Fiji-N5, HDF5, Terafly, and our BioimageVision software were assessed under identical computational conditions. All quantitative analyses were conducted on a dedicated workstation equipped with a CPU (13th Gen Intel® Core™ i9-13900KF utilizing 8 performance cores), RAM (64 GB DDR4 at 2133 MHz), and an operating system (Windows 10). The storage configuration included a Seagate IronWolf Pro 4TB HDD and a WD_BLACK P40 2TB SSD. This setup ensured consistent resource allocation across all tests.

## Acknowledgments

We thank Profs. Gökhan S. Hotamışlıgil, Meng Cui, Scott M. Sternson, and Paul W. Tillberg for making their imaging data available to the public. We thank Prof. Hu Zhao for providing the light-sheet microscopy images. We thank Profs. Jing Yuan and Yu Hua for their valuable suggestions. We also thank the Optical Bioimaging Core Facility of WNLO-HUST for the support in data acquisition.

## Author contributions

Conceptualization: S.Z. and T.Q.

Methodology: T.Q., X.Q., L.C., H.Z., and N.L.

Test Datasets Production: N.L. and X.L.

Software and Code Writing: L.C., X.Q., X.G., S.G., S.L., and J.H.

Data Analysis: T.Q., X.Q., H.Z., X.L., and H.Z.

Writing: T.Q., L.C., X.Q., and N.L.

## Data and materials availability

The test data in Figure 1 is available in https://zenodo.org/records/15054719 and https://zenodo.org/records/14607784. The 100G datasets in Figure 2 is available in https://zenodo.org/records/15173679 and https://zenodo.org/records/15173982. Other datasets in 2, 4 and 5 are extremely large and can be available on request from the corresponding author. Other datasets are publicly available from the previous studies.

All codes, the software tool BioimageVision and its user guide are available on GitHub: https://github.com/Quanlab-Bioimage/BioimageVision. The Dec-Tiff is available on GitHub: https://github.com/Quanlab-Bioimage/Decoupled-Tiff. The compatibility of Bio-VS into Napari and Fiji is available on GitHub: https://github.com/Quanlab-Bioimage/BV-format-compatibility.

