## Supplementary_materials for "Fast Loading and Transformation of Large-Volume Bioimaging Data Consistently Approaching the I/O Peak Speed"

Lin Cai *et al.*

**This PDF file includes:**

Figs. S1 to S7

Table S1

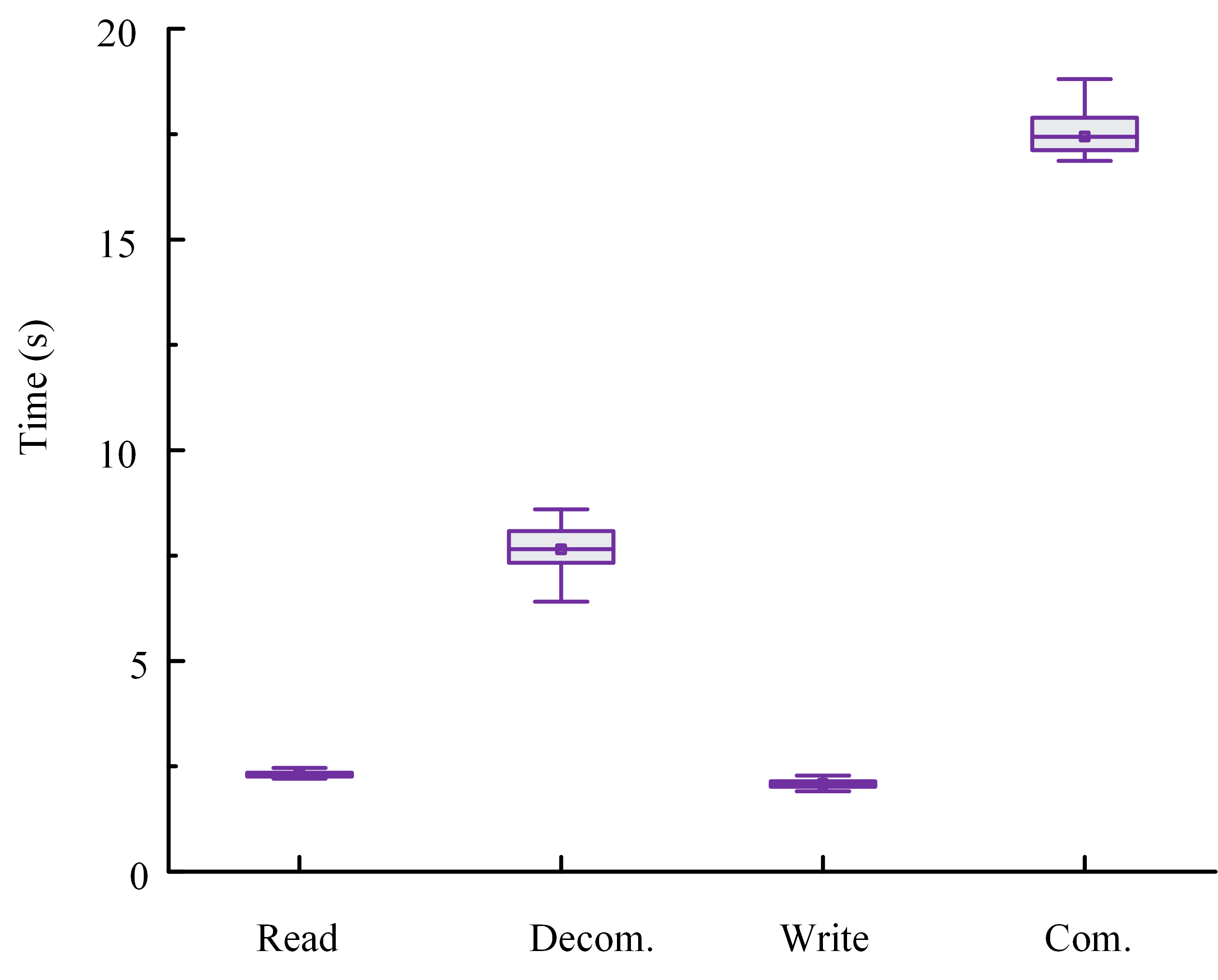

Fig. S1. Time cost of image reading, decompression, writing and compression. The test data is a sequence of 40 images, each with size of 30720×33792. For each image, we directly measured the total time required for both reading and decompression. By subtracting the measured reading time from this total, we derived the decompression time. This approach was applied to determine the time costs associated with image writing and compression.

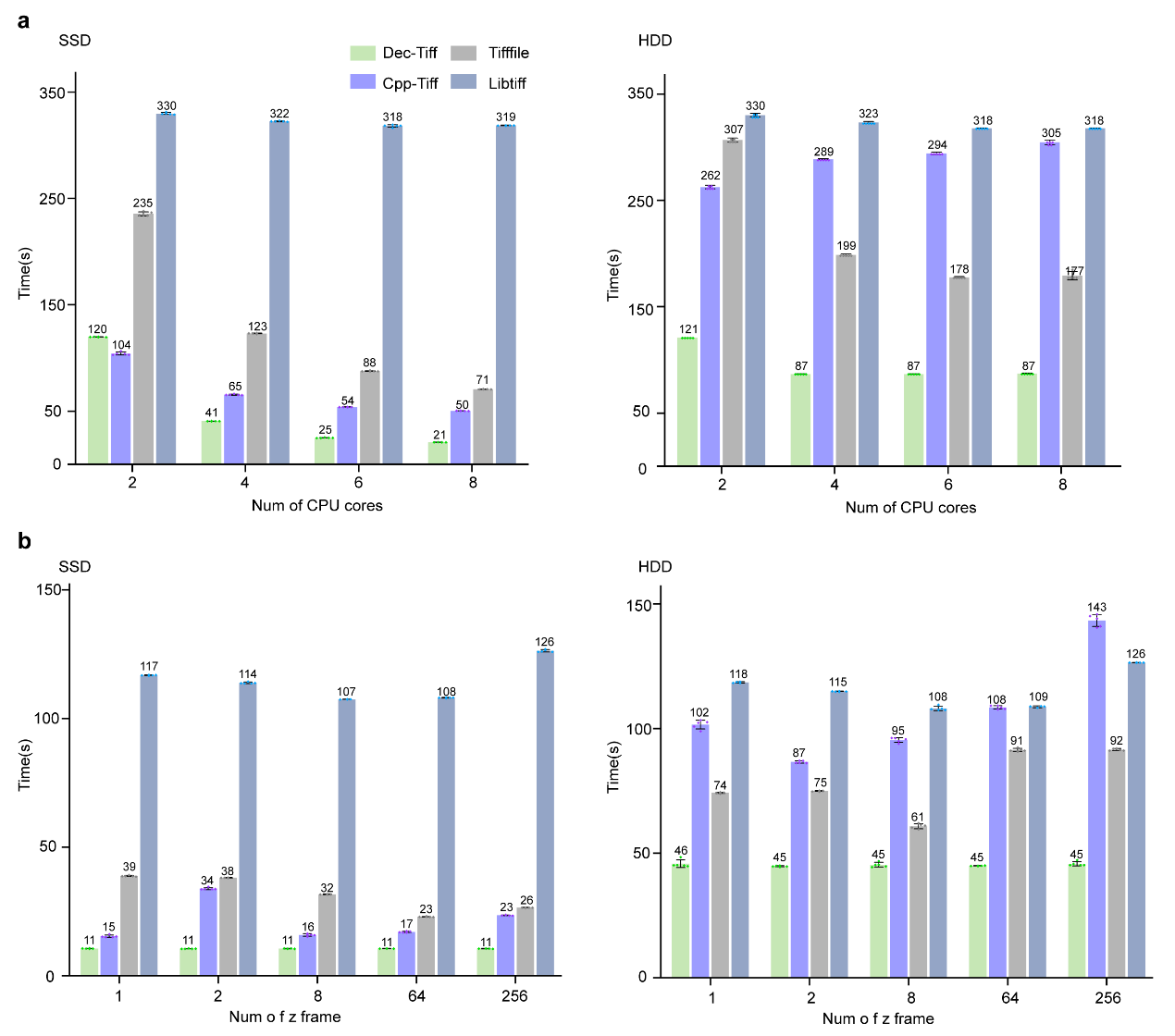

Fig. S2. Quantitative Comparison of image loading performance across diverse conditions. (a) Performance bench-marking of TIFF Libraries on multi-core CPUs. A comparative analysis of the loading times derived from Dec-Tiff, Cpp-Tiff, Tifffile, and Libtiff libraries when processing image stacks of uint16 data type, possessing dimensions of 650×500×256 (x, y, z). This evaluation was performed across varying numbers of CPU cores to assess the scalability and efficiency of each library in multi-threaded environments. (b) Impact of image stack depth on loading times. The influence of image stack depth, specifically the number of z-frames, on the loading performance of the aforementioned libraries. This analysis was conducted using a diverse set of image stacks, all of uint16 data type, but varying in their dimensional configurations: 11000×11000×1, 11000×5500×2, 5000×2500×8, 1200×1350×64, and 650×500×256. By varying the number of z-frames, we aimed to evaluate the robustness of each library when confronted with image data of varying complexities.

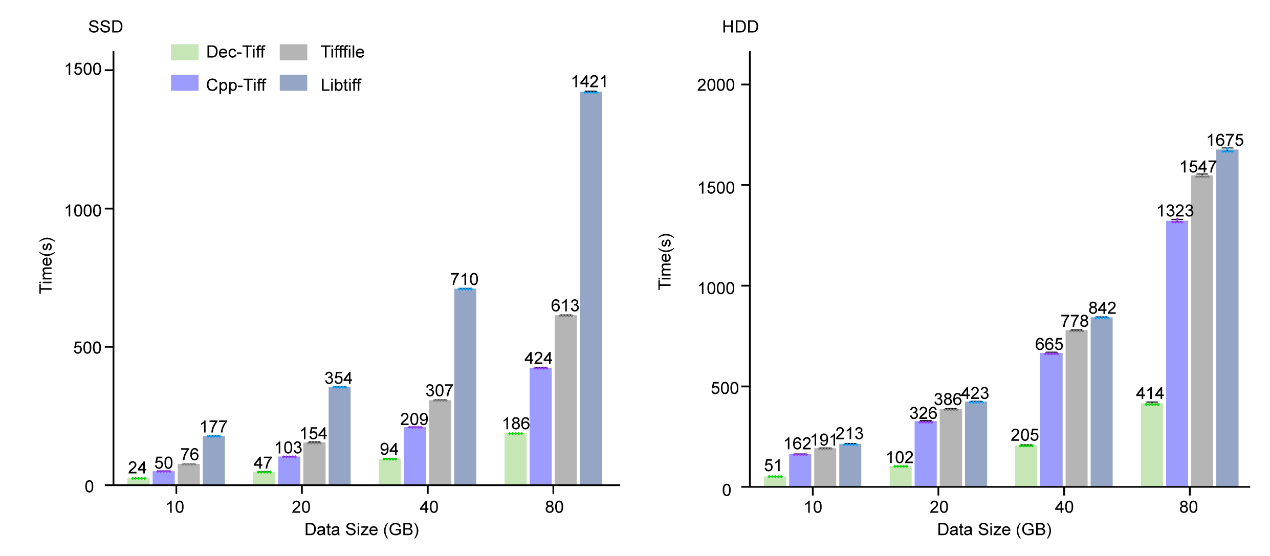

Fig. S3. Comparative evaluation of image saving performance derived from several TIFF libraries. The time required to save image data using Dec-Tiff, Cpp-Tiff, Tifffile, and Libtiff libraries across a range of data sizes. To control the data size s, each library was tasked with saving the same uint16 image block, possessing dimensions of 650×500×256 (x, y, z).

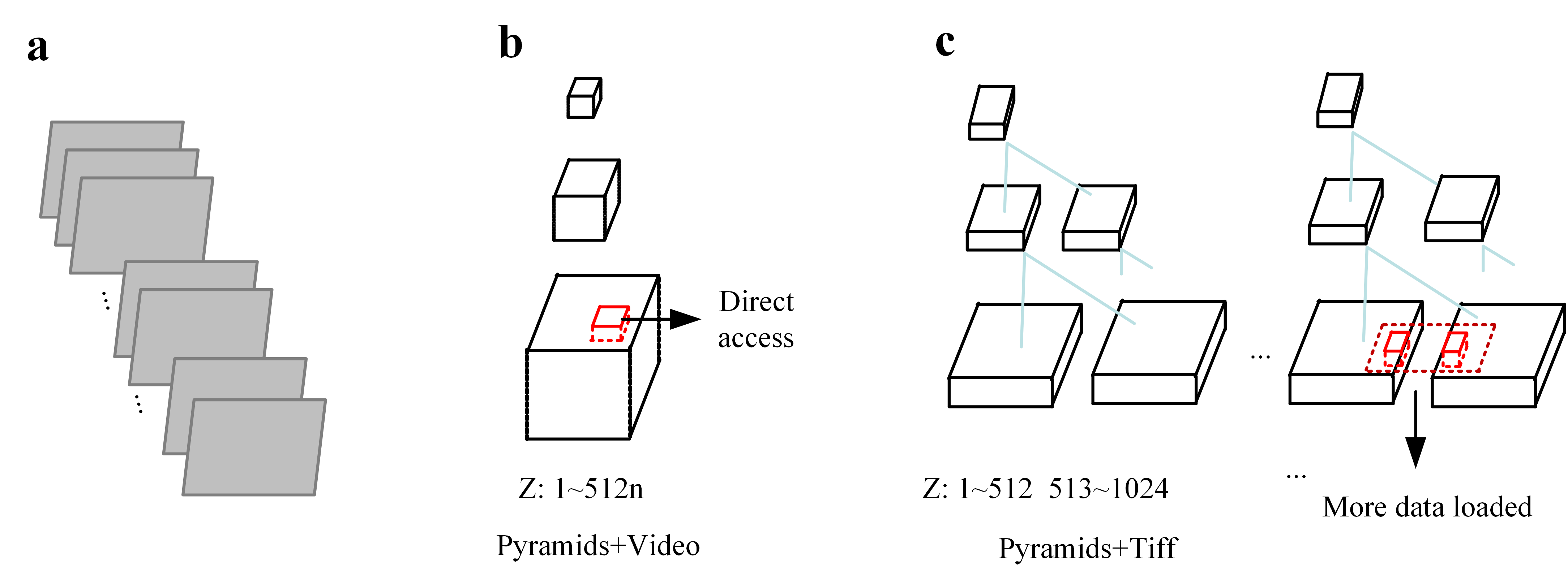

Fig. S4. Comparison of pyramid data structures. (a) An image sequence that requires conversion into a pyramid data structure, a hierarchical organization essential for efficient data management and access. (b) Non-divided z-axis chunk pyramid structure. This pyramid data structure is characterized by chunks that are not subdivided along the z-axis. Notably, these chunks are stored in a video format, which facilitates direct access to specific data regions (as indicated by the red rectangular box). This direct access capability enhances efficiency by minimizing data retrieval overhead. (c) Traditional fixed-length z-axis pyramid structure. In contrast to the non-divided z-axis structure, the traditional pyramid data structure employs fixed-length chunks along the z-axis. These chunks are based on the TIFF format. A limitation of this structure is that accessing a specific block often necessitates loading additional, non-required data (depicted by the dashed rectangular region), potentially increasing data handling inefficiencies.

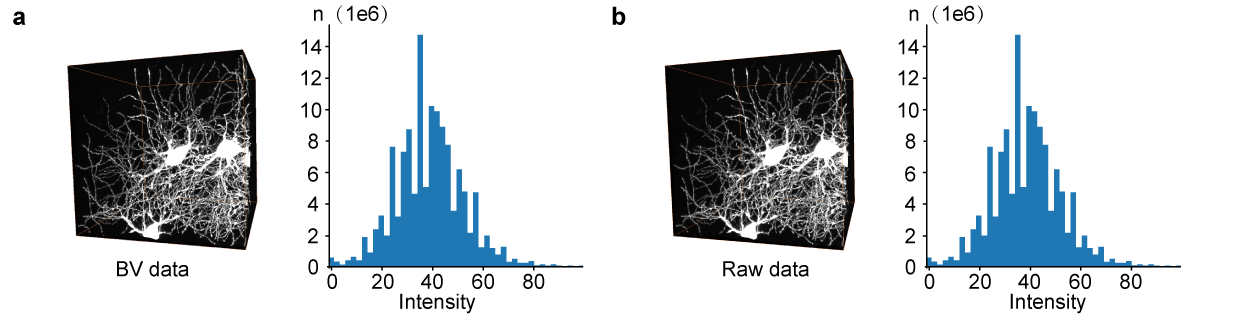

Fig. S5. Validation of lossless compression for BV Data. (a) BV data block and statistical distribution. This panel presents a block of BV data, measuring 512×512×512 in size, accompanied by its corresponding statistical histogram, which illustrates the distribution of data intensities. (b) Raw data block and histogram. The raw data block equivalent to that displayed in (a), also paired with its statistical histogram. To rigorously verify the lossless compression of the BV data, a pixel-wise comparison was performed between the compressed and raw data blocks. The outcome of this comparison is a zero matrix, demonstrating that the compression process has preserved all original data without any loss.

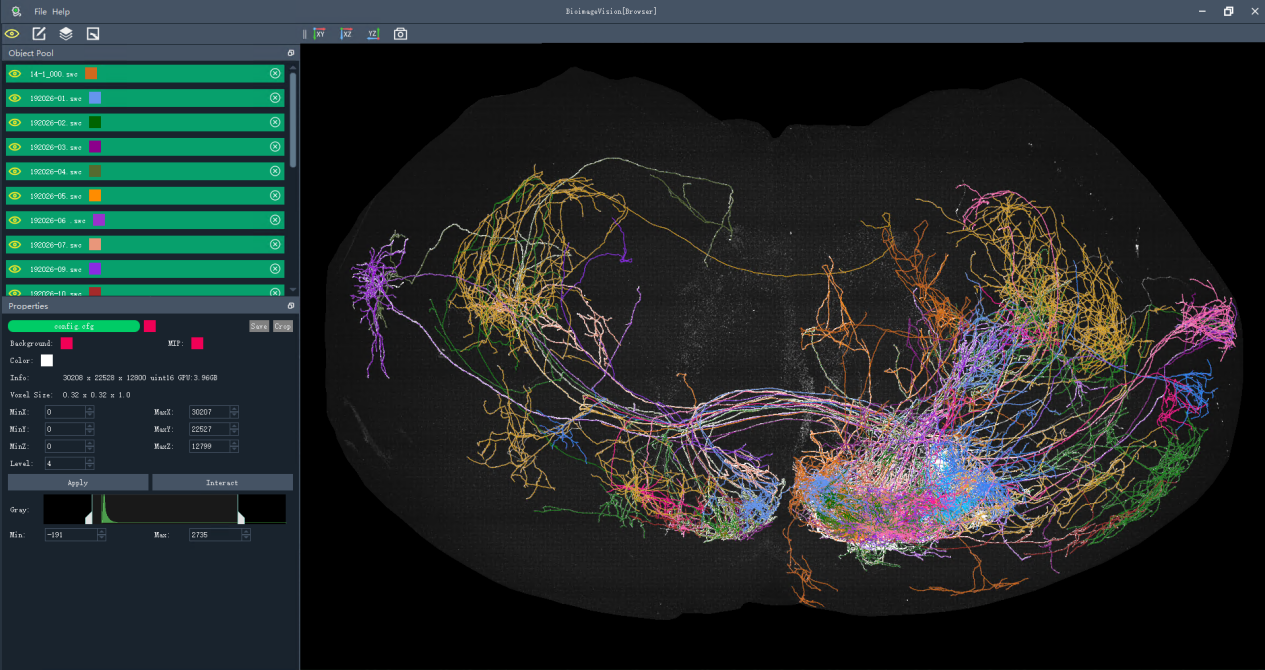

Fig. S6. Visualization of whole mouse brain and reconstructed neurons using BioimageVision. The whole mouse brain data collected with fMOST and its reconstructed neurons are displayed simultaneously. The reconstructed neurons are identified by different colors.

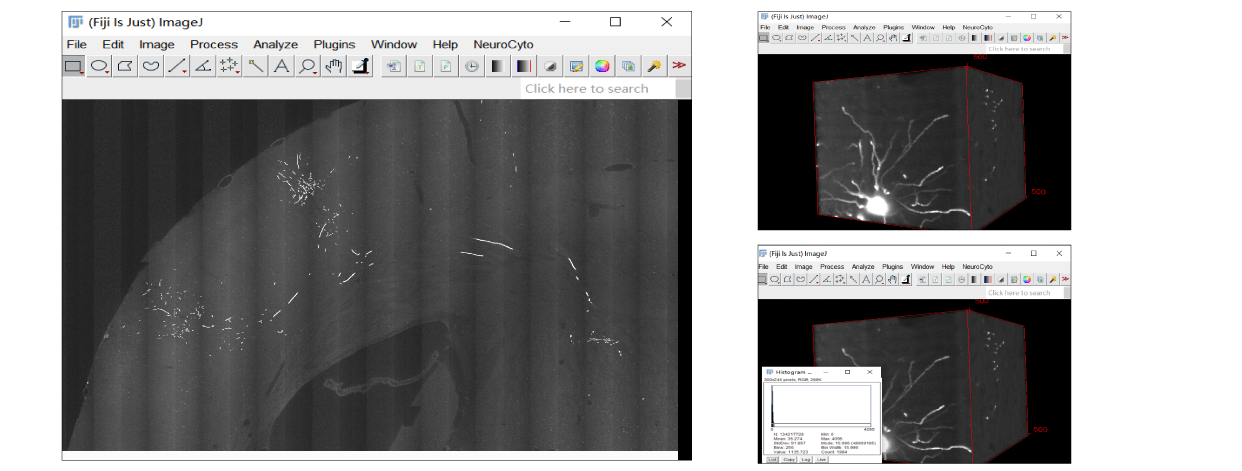

Fig. S7. Integration of Bio-VS data format into Fiji. The first column presents Bio-VS format data that has been imported into Fiji, ready for analysis. The second column illustrates a selected Region of Interest (ROI) and the subsequent calculation of pertinent statistical indicators, all achieved utilizing Fiji’s built-in plugins. This demonstrates the platform's capability to handle and analyze Bio-VS data effectively.

Table. S1. A comparison of the time taken to generate pyramid data, as well as the size of the resultant pyramid data, when utilizing Bio-VS versus Ome-zarr. The testing datasets include four volume images generated with fMOST imaging system, each with size of 100 GB.

| Data Description | Bio-VS  Time (s) | Bio-VS  Size (GB) | Ome-zarr  Time (s) | Ome-zarr  Size (GB) |
| --- | --- | --- | --- | --- |
| Low-SNR data  (10240×12288×500,16-bit) | 242 | 38.8 | 1191 | 44.6 |
| Low-SNR data  (8192×8192×1600,8-bit) | 222 | 30.0 | 1262 | 64.0 |
| High-SNR data  (8192×8192×1600,8-bit) | 147 | 22.3 | 1125 | 30.5 |
| High-SNR data  (8192×8192×1600, 8-bit) | 125 | 19.7 | 1083 | 26.1 |
